# A novel inhibitor of complement C5 provides structural insights into activation

**DOI:** 10.1101/647354

**Authors:** Martin P. Reichhardt, Steven Johnson, Terence Tang, Thomas Morgan, Nchimunya Tebeka, Niko Popitsch, Justin C. Deme, Matthijs M. Jore, Susan M. Lea

**Author notes:** MRC Laboratory of Molecular Biology, Cambridge, UK. Department of Biochemistry, University of Oxford, Oxford, UK. Department of Physiology, Anatomy and Genetics, University of Oxford, Oxford, UK. Institute of Molecular Biotechnology of the Austrian Academy of Sciences (IMBA), VBC, Vienna, Austria.

## Abstract

The complement system is a crucial part of innate immune defences against invading pathogens. The blood-meal of the *tick Rhipicephalus pulchellus* lasts for days, and the tick must therefore rely on inhibitors to counter complement activation. We have identified a novel class of inhibitors from tick saliva, the CirpT family, and generated detailed structural data revealing their mechanism of action. We show direct binding of a CirpT to complement C5 and have determined the structure of the C5-CirpT complex by cryo-electron microscopy. This reveals an interaction with the peripheral macro globulin domain 4 (C5_MG4) of C5. To achieve higher resolution detail, the structure of the C5_MG4-CirpT complex was solved by X-ray crystallography (at 2.7 Å). We thus present the novel fold of the CirpT protein family, and provide detailed mechanistic insights into its inhibitory function. Analysis of the binding interface reveals a novel mechanism of C5 inhibition, and provides information to expand our biological understanding of the activation of C5, and thus the terminal complement pathway.

## Introduction

The bloodmeals of some ticks may last several days, providing ample time for their target to mount a full immune response against the tick during their feeding, and ticks within the same colony will re-bite individuals multiple times, further enhancing immune responses and exposing the tick to their deleterious effects. To survive, ticks have evolved potent inhibitors of mammalian immunity and inflammation. Tick saliva thus represents an interesting target for the discovery of novel immune system modulators, in particular, inhibitors of the very early initiators of inflammation, such as the complement system.

The complement system plays a major role in targeting the innate immune defense system, and is primarily involved in anti-microbial defense, clearance of apoptotic cells and immune complexes, and finally immune regulation (1, 2). Activation may be initiated by target-binding of pattern recognition molecules, such as C1q (classical pathway), mannose binding lectin (MBL), ficolins (1–3) or collectins (10–12) (lectin pathway) (3). In addition, the alternative pathway may auto-activate, including targeting of endogenous surfaces, where inhibitor molecules then terminate further activation (1, 2). The three pathways all converge at the activating cleavage of C3 into C3a and C3b, and the subsequent activating cleavage of C5 into C5a and C5b. C3a and C5a are potent anaphylatoxins acting as soluble inflammatory mediators, while C3b and C5b are deposited on target surfaces. C3b and its inactivated form iC3b function as opsonins for phagocytes, while C5b initiates the terminal pathway by assembly of the pore-forming membrane attack complex (MAC, C5b-C9) (4).

With the ability of complement to target self-surfaces, and induce potent inflammatory responses, the appropriate regulation of complement is essential. Insufficient control of activation is associated with excessive inflammation, tissue damage and autoimmunity (5, 6). Inhibiting activation of C5, and thus the generation of C5a and MAC, has shown great therapeutic benefit in complement-driven inflammatory diseases, such as atypical haemolytic uremic syndrome (aHUS) and paroxysmal nocturnal hemoglobinuria (PNH) (7, 8). The specific targeting of C5 limits the potency of complement activation, while still allowing the effects of the upstream opsonization by C4b and C3b, as well as the immune signaling mediated through C3a. The treatment consists of an anti-C5 antibody that blocks convertase-binding (Eculizumab) (9, 10). However, this antibody is one of the most expensive drugs in the world, and further therapeutic developments are therefore important. Novel inhibitors, such as the tick protein OmCI (Coversin)), the RNAi Aln-CC5, as well as two anti-C5 minibodies (Mubodina and Ergidina) are currently undergoing clinical trials, but a better mechanistic understanding of the activation of C5 is necessary to fundamentally improve the therapies for diseases associated with uncontrolled complement activation (11, 12).

To address the need for a more detailed understanding of the mechanisms of C5 activation, we have identified and characterised a novel family of C5 inhibitors from tick saliva, hereafter named the CirpT (<u>C</u>omplement <u>I</u>nhibitor from *<u>R</u>hipicephalus <u>p</u>ulchellus* of the <u>T</u>erminal Pathway) family. We show that the CirpT family of inhibitors functions by targeting a novel site on C5, and thus provide essential mechanistic evidence for our understanding of C5 activation. We present the cryo-electron microscopy (cryo-EM) structure at 3.5 Å of human C5 in complex with the previously characterised tick inhibitors OmCI and RaCI, as well as a member of the CirpT family (CirpT1). Based on the cryo-EM structure we then solved the 2.7 Å crystal structure of CirpT1 bound to macroglobulin domain 4 of C5 (C5_MG4). Analysis of the specific binding interactions between C5 and CirpT1 suggests that the CirpT family functions by direct steric blocking of the docking of C5 onto the C5-convertase, and our data thus provide support for previous models of C5 activation, which include convertase binding through C5 domains MG4, MG5 and MG7.

## Results

### Identification of a novel complement inhibitor

To identify novel complement inhibitors from tick saliva, salivary glands from the tick *Rhipicephalus pulchellus* were extracted, homogenized, and fractionated utilizing a series of chromatographic methods. In each step, flow-through and elution fractions were tested for their ability to inhibit MAC deposition in a standard complement activation assay using human serum (Wieslab), and the active fractions were further fractionated. The proteins in the final active fraction were analysed by ESI-MS/MS following trypsin digest. Initial analysis against a peptide database generated from the published transcriptome database (13) gave no relevant hits. Therefore, a novel tick sialome cDNA library was assembled from raw sequence data from the Sequence Read Archive (Accession no.: PRJNA170743, NCBI). Using a peptide library generated from this novel transcriptome, we obtained a list of 44 protein hits, out of which twelve contained a predicted N-terminal signal peptide, and were not previously found to be expressed in the published database. These were expressed recombinantly in *Drosophila melanogaster* S2 cells and the culture supernatants tested for complement inhibitory activity. One protein, subsequently termed <u>C</u>omplement <u>I</u>nhibitor from *<u>R</u>hipicephalus <u>p</u>ulchellus* of the <u>T</u>erminal Pathway (CirpT), was shown to inhibit MAC assembly regardless of the initiation pathway of complement (Figure 1a). To identify potential biologically relevant homologues, the CirpT sequence was used to query the expressed sequence tag (EST) database (NCBI) as well as in-house *R. appendiculatus* and *R. pulchellus* sialomes. This search revealed that the protein is highly conserved among ticks, with homologues found throughout the genii *Rhipicephalus* and *Amblyomma*, as well as in the species *Dermacentor andersonii* and *Hyalomma marginatum* (Figure 1c). The proteins identified fall into four distinct clusters with sequence identity varying between 62.6% and 88.3% within the clusters. For further investigation of the mechanisms of these novel types of complement inhibitors, a member from each cluster (hereafter named CirpT1-4) was expressed in *D. melanogaster* S2 cells and all were shown to inhibit complement activation (Figure 1b). For subsequent studies, all four homologues were expressed in *E. coli* SHuffle cells with an N-terminal His-tag and purified using Ni-chelate chromatography, ion-exchange and size exclusion chromatography.

**Figure 1:**
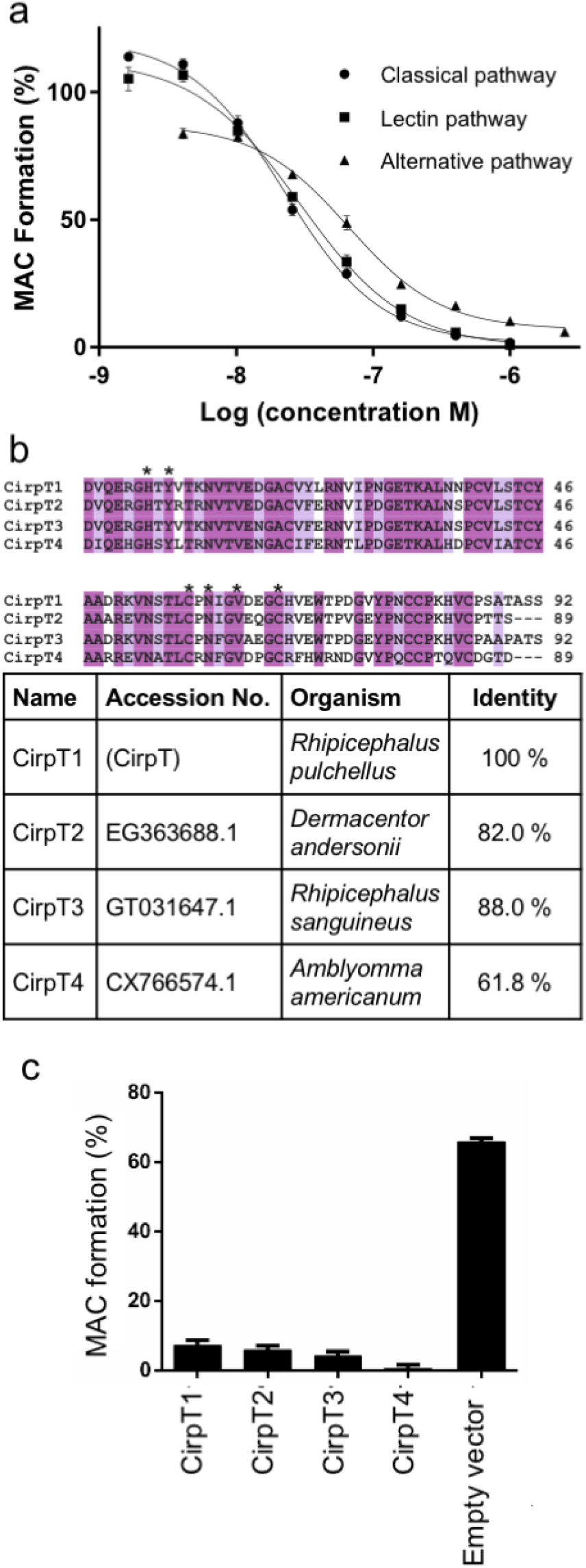
A novel class of tick proteins (CirpTs) inhibit complement activation. Sequential fractionation of tick salivary glands revealed a novel family of tick complement inhibitors. Following recombinant expression, purified tick proteins and culture supernatants from S2-insect cells over-expressing tick molecules were utilized to determine the potential for complement inhibition. **a**) In the commercial Wieslab assay, inhibition of pathway-specific complement activation was tested. Addition of CirpT1 purified from *E. coli* reveal a dose-dependent inhibition of MAC deposition in all three pathways. The lines are showing non-linear fits. Error bars: standard error of mean (SEM), n = 3. **b**) Clustal Omega (EMBL-EBI) sequence alignment of CirpT1-4. The native signal peptides are omitted from this alignment. Colouring based on sequence identity. Stars denote residues relevant for protein interaction (see below). Sequence identity to CirpT1 is indicated in the table. **c**) In the commercial Wieslab Alternative pathway assay, ELISA-wells are coated with LPS and will lead to complement activation and deposition of the membrane attack complex (MAC). Addition of culture-supernatants from insect cells expressing CirpT1-4 inhibit MAC-formation. No inhibition is seen with supernatants from cells transfected with an empty vector, error bars: SEM, n = 3.

All three complement activation pathways were affected by CirpT inhibition, yet no impact was observed on C3 activation (data not shown). The target of CirpT was thus likely to be in the terminal pathway of complement. To pinpoint the specific ligand of CirpT, a pull-down assay from human serum was performed utilizing CirpT1 that was covalently coupled to beads. This identified C5 as the target of CirpT inhibition (Figure 2a). Following this, binding of each CirpT1-4 to C5 was assayed by surface plasmon resonance (SPR). C5 was coupled onto a CM5 chip surface by standard amine-coupling, and CirpT1-4 were flown over in varying concentrations. Analysis of the binding curves for CirpT1 using a 1:1 kinetic model yielded a dissociation constant of 10 × 10^−9^ M (averaged over three single experiments with varying amounts of C5 coupled to the chip surface). Attempts to produce data with reliable fits for CirpT2-4 were unsuccessful, in part due to long dissociation times and an inability to regenerate the binding surface without denaturing the C5. However, the magnitude of binding signal at comparable concentrations demonstrates that they likely bind tighter than CirpT1, with CirpT4 displaying an especially slow dissociation rate. The K_D_ of all CirpT-C5 affinities are thus in the range of 10 nM or tighter.

**Figure 2:**
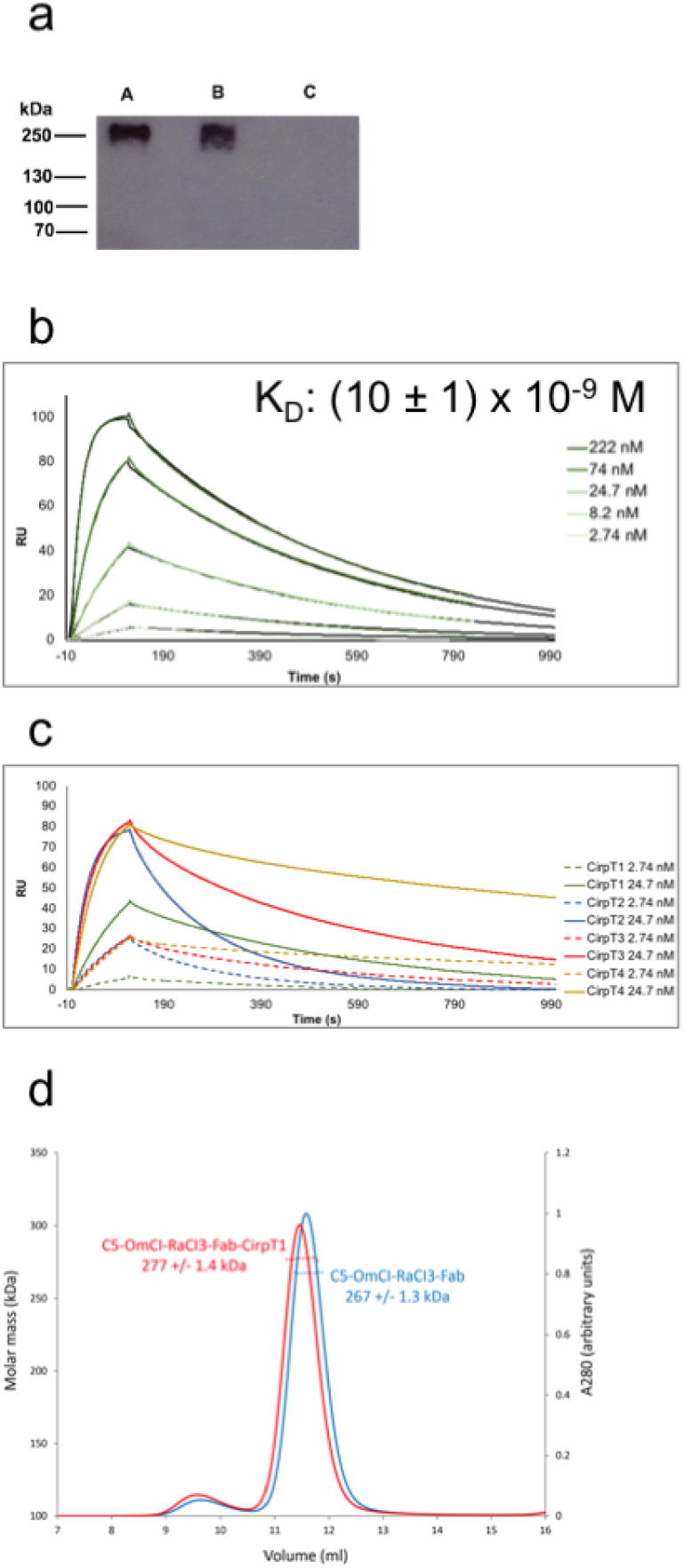
CirpT1 binds C5 through a novel inhibitory binding site. **a)** Western-blotting of serum pull-down. 0.5 mg/ml purified CirpT1 was immobilized on NHS-activated magnetic beads (Pierce, Thermofisher Scientific) and incubated with human serum. Eluted proteins were separated by 4-12 % gradient SDS-PAGE and visualized by Western blotting using a polyclonal anti-C5 antibody. Pull-down lanes A: CirpT1, B: OmCI, C: beads only. **b)** Surface plasmon resonance performed with purified C5 coupled to a CM5 chip by amine-coupling. CirpT1 was flown over in a concentration series from 2.74 – 222 nM, as indicated. Shown are representative curves of CirpT1 flown over surface with three different levels of coupled C5. An approximate dissociation constant was calculated by kinetic curve-fitting using the BiaEvaluation software package (n = 1). **c)** Surface plasmon resonance of CirpT1-4. All CirpTs clearly bind. The binding curves of CirpT2-4 could not be reliably fit, but our data show even tighter interactions for CirpT2-4 than CirpT1. **d)** SEC-MALS traces of purified C5 complexed with the inhibitory molecules OmCI, RaCI3, CirpT1 and the Fab-fragment of the commercial antibody Eculizumab. Binding of CirpT1 does not compete with any of the other inhibitory molecules, revealing a novel mechanism of inhibition.

It was previously demonstrated that other tick-inhibitors targeting C5, OmCI, RaCI, SSL7 as well as the Fab-fragment of the commercially available C5-inhibitory antibody, Eculizumab, have different binding sites on C5 (9, 14). We next sought to compare the binding mechanism of the CirpTs to the previously known modes of inhibition. C5, OmCI and RaCI were purified and complexed with the Eculizumab Fab-fragment and the mass was determined by SEC-MALS. A 10 kDa increase in the mass of the complex was observed when CirpT1 was added to the quaternary complex demonstrating that the binding site for CirpT does not overlap with any previously known inhibitors. The mass increase was consistent with CirpT binding as a monomer. CirpT thus has a novel mechanism for inhibition of complement C5 activation.

### Cryo-EM structure of the C5-OmCI-RaCI-CirpT1 complex

To characterize the novel mechanism of C5 inhibition, we next identified the binding site of CirpT1 on C5 by cryo-electron microscopy. As OmCI and RaCI lock C5 into a less flexible conformation, our approach targeted the full C5-OmCI-RaCI-CirpT1 complex. Purified OmCI-C5 was incubated with a 2-fold molar excess of RaCI and CirpT1, and the complex was purified by size exclusion chromatography. The complex was imaged on a Titan Krios and a 3D reconstruction was generated at a nominal overall resolution of 3.5 Å from 118,365 particles (Figure 3a). The volume generated allowed us to dock and refine the previously determined crystal structure of the C5, OmCI and RaCI1 complex. As observed in the ternary complex crystal structures, local resolution varied across C5 within the complex, with the C345c domain being the least well-ordered part of the complex (5-6 Å). However, domains buried within the core of the C5 demonstrated features consistent with the experimentally determined local resolution of 3.35 Å, including detailed sidechain density (Figure 3b). By comparing the expected density from the crystal structure of the C5-OmCI-RaCI-complex to our newly generated map, we identified an extra density in the lower right corner of the complex, which we attributed to CirpT1. The local resolution in this area was worse than for the overall complex (4-5 Å), indicating a higher level of flexibility. The resolution of the map corresponding to CirpT1 was insufficient to build a *de novo* atomic model of CirpT1. However, the residual density suggested that the main interaction between C5 and CirpT1 was mediated through binding to C5_MG4 (C5 residues L349 - S458).

**Figure 3:**
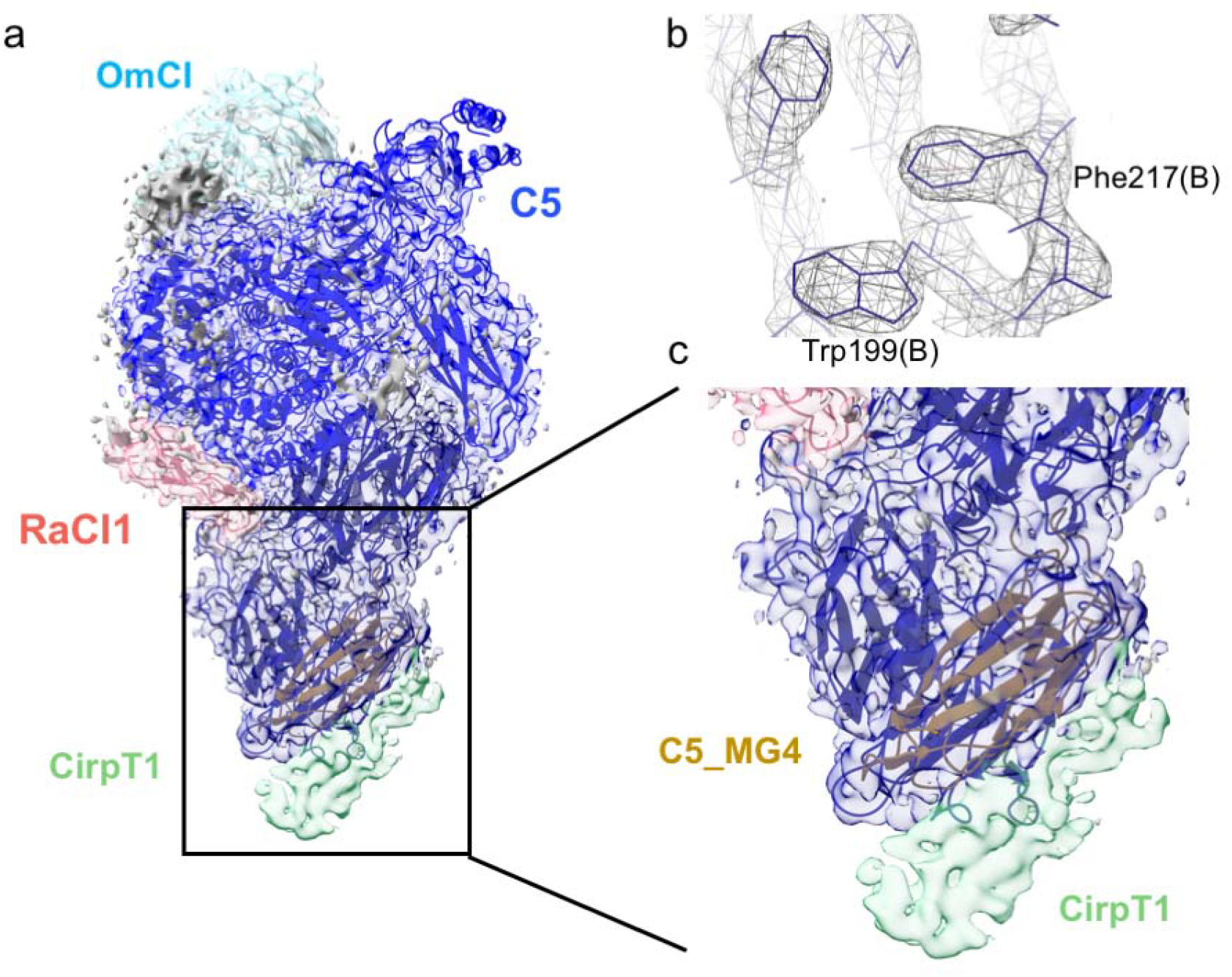
Cryo-electron microscopy structure of the C5-OmCI-RaCI-CirpT1 complex. **a**) Side view of the density map with C5 (blue), OmCI (light blue) and RaCI1 (red) structures built in. Residual density (green) was observed attached to the macroglobulin domain 4 (C5_MG4, gold). This residual density was attributed to CirpT1. **b**) Example of higher resolution data in the density map, allowing the placement of amino acid side chains in the density. The map represented here has been postprocessed with B-factor sharpening **c**) Zoom of C5_MG4 with the CirpT1 density clearly visible. The map represented here is filtered specifically to provide best detailed information for CirpT1.

To understand the interaction between CirpT1 and C5 in greater detail, a complex of CirpT1 and C5_MG4 was crystallized. To this end, the C5_MG4 domain was cloned with an N-terminal His-tag, expressed in *E. coli*, and purified by Ni-chelate and size-exclusion chromatography. To confirm our previous observation of a direct interaction between CirpT1 and C5_MG4, binding was tested by co-elution on a size exclusion column. This yielded a C5_MG4-CirpT1 complex, which was purified, concentrated and crystallized. X-ray diffraction data to 2.7 Å were collected at the Diamond Light Source (beamline I03). The structure of the C5_MG4-CirpT1 complex was solved by molecular replacement (MR) with the isolated C5_MG4 extracted from the C5-OmCI-RacI complex. Initial phases from the partial MR solution produced a map into which a model of CirpT1 was built *de novo*, and the complex refined to give the model described in Table 1 (Figure 4).

**Figure 4:**
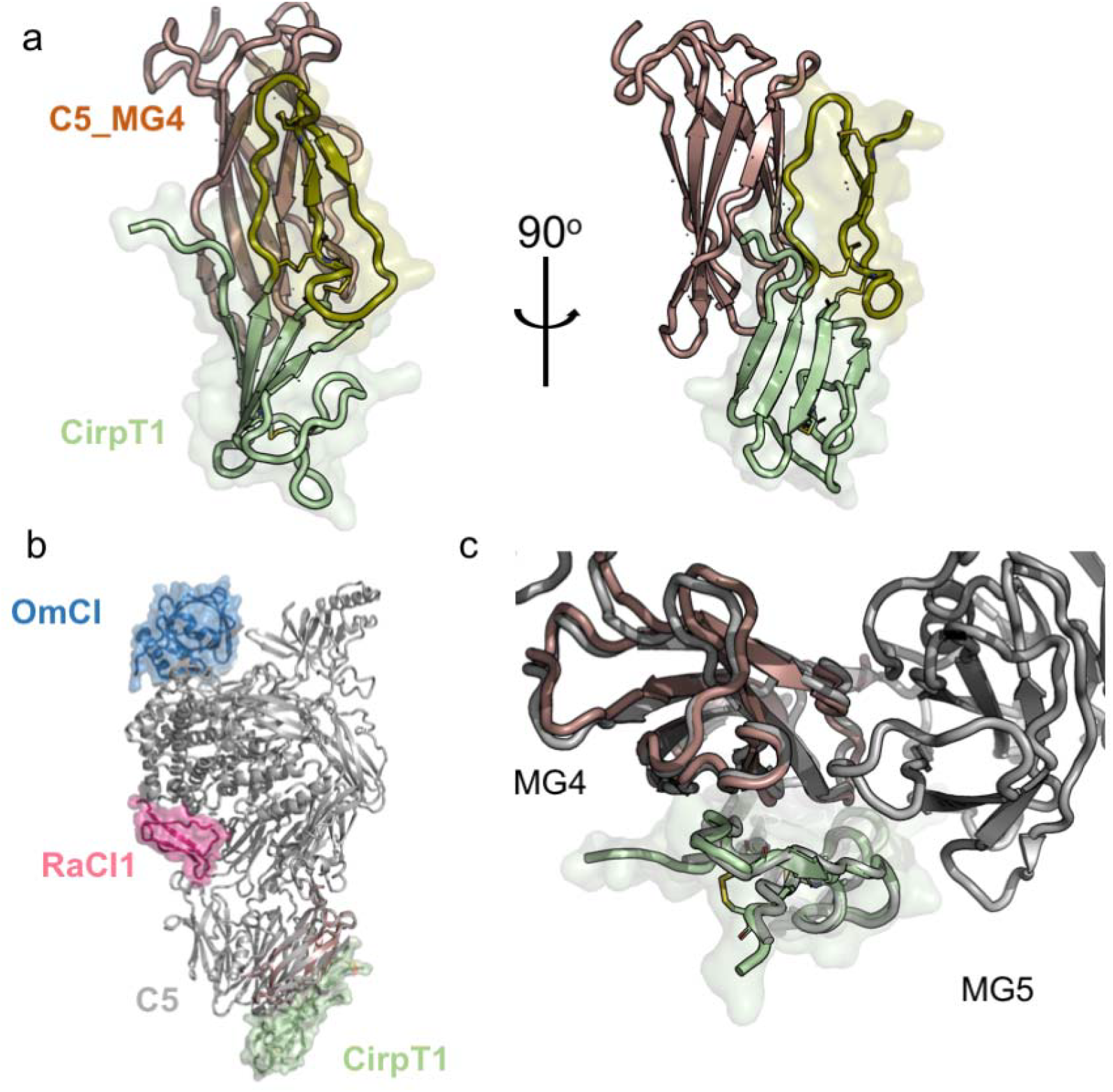
Crystal structure of the C5_MG4-CirpT1 complex at 2.7 Å.. **a**) Front and side views of the crystal structure of the C5_MG4-CirpT1 complex. C5_MG4 shown in brown, CirpT1 shown in green, with the bulky subdomain highlighted in pale green, and the flat subdomain in olive green. Disulphide bridges are shown in yellow. **b**) Overlay of the C5_MG4-CirpT1 structure with the full C5-OmCI-RaCI1-CirpT1 complex reveal that CirpT1 sits in the density observed from the cryo-EM (front view). C5 shown in grey, RacI1 in red and OmCI in blue. **c**) A closer investigation of the placement of CirpT1 shows it sits between C5_MG4 and C5_MG5 (Top view). Though the major interaction is with C5_MG4, and this is sufficient for binding, the structural overlay shows a potential for interaction with the C5_MG5 as well.

CirpT1 is made up of two domains; a bulky N-terminal domain, and a flat looped C-terminal domain (Figure 4). An FFAS search (15)of the CirpT1 sequence showed overall similarities to the von Willebrand factor type C (VWC) domain family, with four conserved cysteine bridges (score: −22.2, 26 % identity). The N-terminus of CirpT1 extends further, however, and engages in a four-stranded β-sheet. Using the Dali server (16), searches for either the bulky N-terminal domain of CirpT1 (aa 2-56) or the full CirpT1 (2–87) identified the porcine beta-microseminoprotein (MSMB, pdbid: 2iz4-a) as the closest structural homologue (r.m.s.d. of 1.8 Å with 46 out of 91 residues aligned for the N-terminal domain, and r.m.s.d. of 4.2 Å with 72 out of 91 residues aligned for the full CirpT1). In both CirpT1 and MSMB, the N-termini form a Greek key motif with four antiparallel strands, with an extended loop between the first and fourth strands. This extended loop folds back under the β-sheet, and is locked into this confirmation by a disulfide-bridge to the third β-strand in the sheet. The C-terminal domain of CirpT1 corresponds more closely to the VWC fold, but a Dali search for this domain alone did not yield any hits. The orientation of the N-terminal and C-terminal domains in relation to each other differ from known homologues, an observation consistent with flexibility reported in ths hinge-region of the von Willebrand folds (15).

A detailed analysis of the binding interface was carried out using PDBePISA (17). This revealed an extensive interface that encompasses most of the C-terminal domain of the CirpT1, with additional contributions from the N-terminal strand and two residues at the tip of the extended loop in the N-terminal domain (Tyr23 and Leu24). The interface is predominantly hydrophobic in nature, with His7, Tyr23 and Asn59 on CirpT1 contributing sidechain hydrogen bonds (to Asp405, Ser426 and Asn423 respectively). (Figure 5). Overlay of the C5_MG4-CirpT1 structure onto the C5-OmCI-RaCI-CirpT1 structure revealed minor clashes between CirpT1 and the neighboring C5_MG5 domain. However, analysis of the cryo-EM volume in this region revealed a small rigid body movement of the CirpT1 away from C5_MG5, resolving the clash. This placement of CirpT1 in the context of intact C5 revealed additional contacts between CirpT1 and C5_MG5, including the creation of a hydrophobic cleft between C5_MG4 and C5_MG5 that Trp70 and Pro72 of CirpT1 pack into. Analysis of the sequence conservation of the CirpTs (Figure 1b) revealed that the residues involved in binding C5 are highly conserved, with residues His7, Tyr9, Cys57, Asn59, Val62 and Cys66 conserved among all four CirpTs examined. Of note, the stretch of residues from Cys66-Glu69 in CirpT1 extend over an extension of the hydrophobic crevice between C5_MG4 and C5_MG5, but leaving a hydrophobic hole. CirpT4, which appeared to have the slowest dissociation rate in the SPR studies, has a Val68Phe substitution which would fill this cavity, further stabilizing the complex.

**Figure 5:**
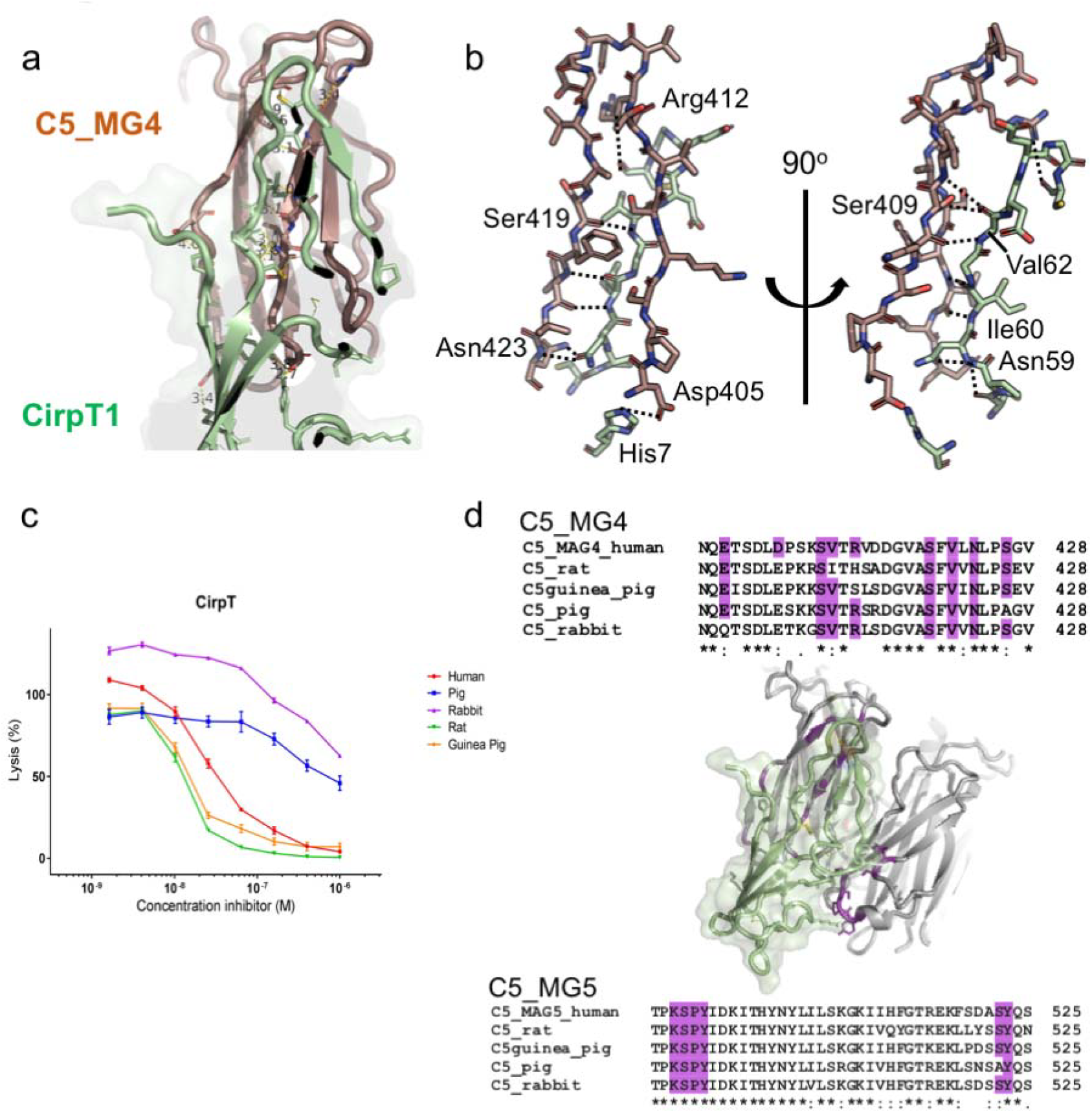
The C5-CirpT1 binding interface. The main binding interface between CirpT1 and C5_MG4 is created by one of the C-terminal strands binding between two β-strands in the C5_MG4 β-barrel. Additionally, two salt-bridges connect the lower loops of the C5_MG4 β-barrel to the N-terminal domain of CirpT1. Conservation of the CirpT1 binding interface on C5 across mammalian species correlates to functional inhibition. **a**) Front view cartoon of CirpT1 (green) binding to C5_MG4 (brown). Interacting amino acids are displayed as sticks. **b**) Close-up back and side views of the loop extension in CirpT1 placed between the two β-strands of C5. Amino acids are shown as sticks. Hydrogen-bonds with atomic distances below 4.0 Å are highlighted in dashed lines. **c**) Serum complement-mediated lysis of red blood cells was assayed with increasing concentrations of CirpT1 inhibitor. A dose-dependent inhibition was observed for all tested species; human, pig, rabbit, rat, and guinea pig. Error bars: SEM, n = 3. **d**) Sequence alignment of C5_MG4 and C5_MG5 of tested species in the PISA-predicted CirpT binding interface. This reveals a high level of sequence conservation, thus explaining the potent inhibitory effect of CirpT across species (residues involved in binding are highlighted in purple).

Ticks feed on a wide variety of mammalian species, and they are therefore required to have potent inhibitors of complement from multiple species. We tested the ability of CirpT1 to inhibit complement mediated red blood cell lysis by serum from rat, pig, guinea pig and rabbit (Figure 6). We observed inhibition in all species investigated, although inhibition by rabbit and pig serum was less potent. Mapping C5 sequence conservation in the binding interface demonstrated that the key residues involved in binding CirpT1 are highly conserved across these species, explaining this broad spectrum of activity. Specifically, the C5-residues Glu398, Ser407, Val408, Ser69, Val419, Asn421 and Ser424, are all highly conserved with only single substitutions in particular species (rabbit Glu398Gln, rat Val408Ile, and pig Ser417Ala). The salt-bridge-forming Asp403 in human is substituted with a glutamic acid in all other species. Arg410 is replaced by a histidine and a serine in rat and guinea pig respectively. In addition, the potential binding interface to C5_MG5 is also highly conserved. LysSerProTyr489-492 and Tyr523 are conserved in all tested species, while Ser522 is found in all species except the pig (substitution to Ala522).

**Figure 6:**
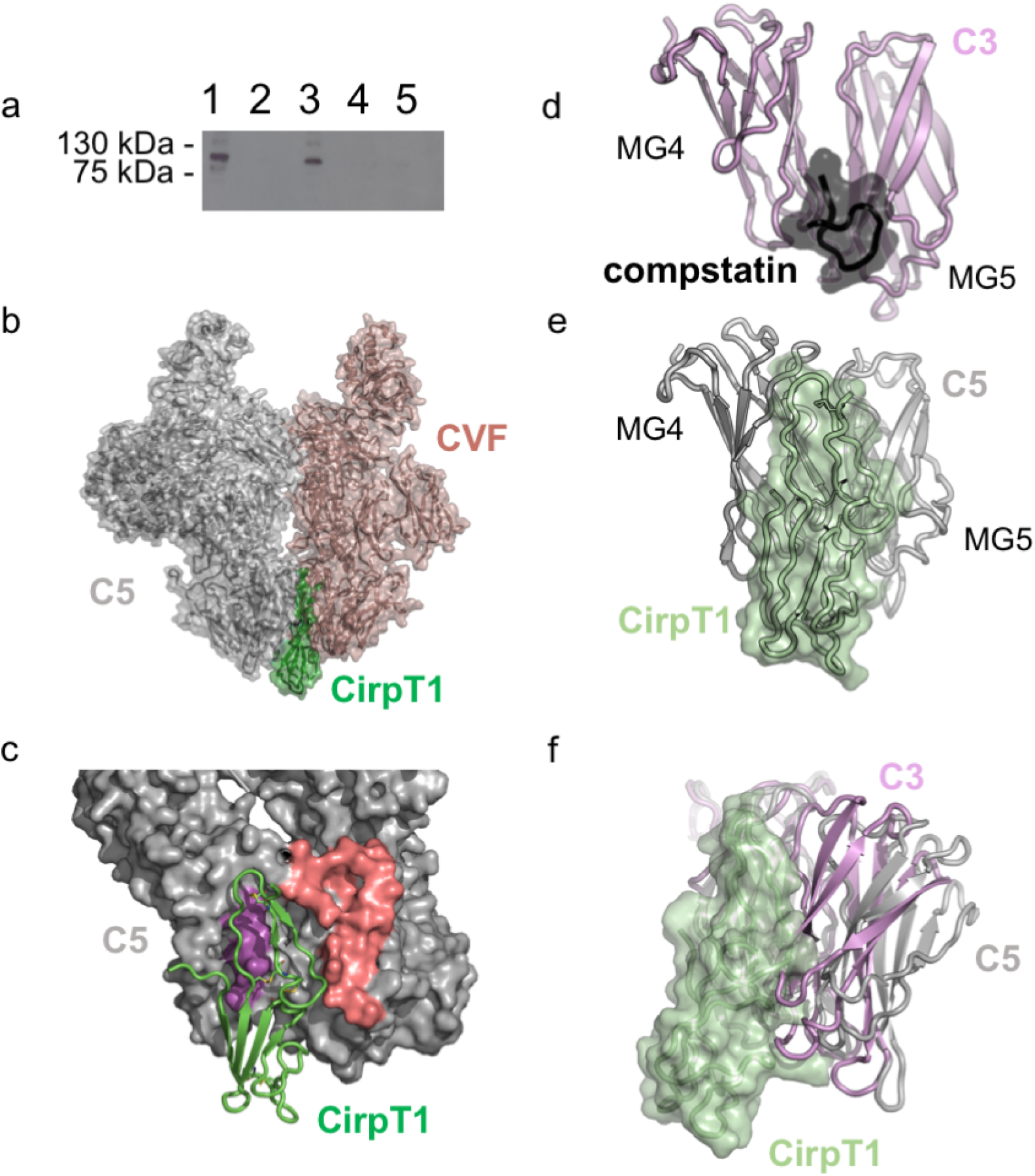
Mechanism of inhibition. The CirpT-family of inhibitors likely mediate their effect by blocking of C5-binding to its convertase. Previous structural data of the C3b homolog, CVF, in complex with C5, suggest the MG4 and MG5 domains are essential for convertase-docking. Our data provide further support to this model of convertase binding. **a**) Western-blotting (reducing conditions) of elutions from C3b-coated magnetic beads. C5 binds to the C3b-coated beads (lane 3), however, this binding is interrupted by addition of CirpT1 (lane 5). Lane 1: Purified C5. Lane 2: C5 + empty beads. Lane 4: C5 + CirpT1 + empty beads. **b**) CirpT (green) overlaid with the complex of C5 (grey) and CVF (brown). CirpT1 sits right at the CVF-C5 binding interface. **c**) Surface representation of C5 with residues essential for CVF binding highlighted in MG4 (purple) and MG5 (red). CirpT binding directly overlaps with the binding site on C5. **d**) Front view of compstatin (black) binding to C3 (pink). **e**) Front view of CirpT1 (green) binding to C5 (grey). These structures reveal that CirpT1 and compstatin block corresponding sites in C5 and C3, respectively. **f**) Sideview of the overlay of the CirpT1 (green) with C5 (grey) and C3 (pink) (aligned by the MG4 domains). The domain organization of C3_MG5 in relation to C3_MG4 shows a much closer conformation as compared to C5. C3_MG5 is thus likely to provide steric hindrance of the CirpT inhibitor, thus explaining the specific C5-targeting of the tick molecule.

### Mechanism of complement inhibition

Current models propose that C5 is activated by docking to C3b, complexed with either of the activating proteases C2a or Bb. This results in C5 cleavage into C5a and C5b. The key interaction between C5 and C3b is thought to involve packing of the MG4 edge of C5 against the C3b-containing convertase, implying that steric block of this event might explain inhibition by the CirpT family. In an effort to deduce the specific mechanism of inhibition, we analyzed the binding of C5 to beads coated with C3b, mimicking the high critical surface concentration of C3b required for C5 conversion, in the presence of CirpT1 (Figure 6).

Based on the binding site of CirpT1 to C5, we predicted that CirpT1 would function by blocking an interaction between C5 and surface-bound C3b. Purified C3 was activated by trypsin-cleavage; the resulting C3b was biotinylated and coupled to streptavidin magnetic beads in a concentration sufficient for C5-binding (30 μg/ml). When CirpT1 was co-purified with C5 prior to incubation with the C3b-coated beads, binding was abolished.

## Discussion

In an effort to understand the mechanisms underlying complement activation, we utilized the complement inhibitory properties of tick saliva. We here describe a novel class of tick inhibitory molecules targeting the terminal pathway of complement: <u>C</u>omplement <u>I</u>nhibitor from *<u>R</u>hipicephalus <u>p</u>ulchellus* of the <u>T</u>erminal Pathway (CirpT). Homologous sequences were identified in several tick species and genii, and fell into four distinct clusters, CirpT1-4. They all share a high degree of sequence identity with CirpT1 (62.6 – 88.3 %), and all inhibit complement activation through binding to C5 with nano-molar affinities. To understand this novel mechanism of complement inhibition, we generated a cryo-electron microscopy structure of C5 complexed with the tick inhibitors OmCI, RaCI and CirpT1. This verified previous crystallographic binding mechanisms of OmCI and RaCI. Furthermore, the generation of a cryo-EM map showing density of CirpT1 allowed a targeted crystallographic approach, revealing the interaction between CirpT1 and the MG4 domain of C5 at 2.7 Å resolution.

The CirpT family adopts an extended two domain fold stabilized by multiple disulphide bonds. An extended strand connects the beta-sheet N-terminal subdomain to the flatter loopy C-terminal domain. This flat domain is further connected to the bulky domain by a disulfide bridge. The limited connection between the two subdomains allow for a flexibility of this “hinge-region”, which can likely account for the small variation between the crystal structure and the cryo-EM structure of CirpT1. Flexibility between the two subdomains is consistent with similar flexibility reported for the von Willebrand folds (15).

Cleavage of the homologous molecules C3 and C5 is carried out by mechanistically similar convertases. The C3 convertase consists of a docking-molecule (C3b or C4b) associated with an activated serine protease (either factor Bb or C2a, respectively). The C3 convertases C4b2a (classical/lectin pathways) and C3bBb (alternative pathway) generate novel surface-associated C3b (hereafter termed C3b’), which subsequently associates with additional factor Bb, and thus provides a potent amplification loop. Following the deposition of critically high concentrations of C3b’ on a surface, a shift in convertase activity is created, permitting C5 as a substrate (18–22). Current models of C5 activation include docking of C5 to C3b, complexed with either of the activating proteases C2a or Bb (9, 22). Previous work utilizing the C3b homolog cobra venom factor (CVF) has provided structural information of how this docking may occur (14). Structural data of C5 complexed with CVF (PDB ID: 3PVM) show that C5 domains MG4, MG5, and MG7 are involved in this interaction. The main binding interface is found with surface-residues of MG4 and MG5. The C5 MG4 and MG5 residues Ser419– Pro425, Thr470–Ile485, and Asp520–Asn527 are all located in proximity to the CVF residues Ser386–Thr389, Ile399–Leu404, Thr450–Lys467, and Arg498–Asn507 (CVF domains MG4 and MG5). The CirpT1 binding interface with C5 directly overlaps with this binding-interface. We therefore propose that the CirpT mechanism of inhibition is through direct steric hindrance of C5-docking to its convertase. This is substantiated by the observed CirpT1-mediated inhibition of C5 binding to surface-coated C3b. Our structural data thus support the CVF-generated model of C5-convertase docking, and provides further evidence for this being a crucial part of the mechanism of C5-activation in humans (Figure 6B and C). C5-convertase activity is dependent on a high density of surface-associated C3b (22, 23). Combined with structural knowledge of C5-inhibitors such as OmCI and RaCI, a novel hypothesis suggesting two distinct binding interfaces between C5 and C3b has emerged. (9, 22, 23). The data presented here indicate, that the binding-interface utilizing C5_MG4 is essential for sufficient binding of C5 to C3b.

Due to the high levels of homology between C3, C5, and their convertases, we compared our proposed mechanism of inhibition to that of a known C3-inhibitor, compstatin (Figure 6D). The structure of compstatin, crystallized at the interface between two C3c molecules (PDB ID: 2QKI), suggesting that inhibition is mediated through steric hindrance of C3 docking onto the C3-convertase (24). An overlay of the C3-compstatin complex with our C5-CirpT1 complex reveals that CirpT1 and compstatin block a similar site on the MG4 domain of either complement molecule. While inhibition is mediated through very similar mechanisms by CirpT1 and compstatin, targeting corresponding areas of C5 and C3, respectively, the binding is specific for each molecule. No binding to or inhibitory effect of CirpT1 has been observed for C3. Despite the lack of amino acid sequence conservation between C3 and C5 at the major binding interface, the placement of the backbone is consistent, and would be expected to mediate binding in both C3 and C5. However, overlay with the molecular structure of C3 (PDB ID: 2a73) reveals that the neighboring C3_MG5 domain packs much closer to C3_MG4, essentially sterically hindering binding of CirpT1 (Figure 6E).

In conclusion, we here present the structural fold of a novel family of complement inhibitors. By identifying the specific binding site, we provide further mechanistic insight into current models of C5 activation. Combined with the detailed molecular understanding of multiple C5 inhibitors, our mechanistic understanding may allow for future developments of clinically relevant therapeutic strategies.

## Methods

### Fractionation of *R. pulchellus* salivary glands

*R. pulchellus* ticks were reared and 250 salivary glands were dissected according to Tan and colleagues (25). The gland protein extract was topped up with 25 mM Na2HPO4/NaH2PO4, pH 7.0 to 10 mL. The sample was then fractionated by sequential anion exchange, reverse-phase hydrophobic interaction and size exclusion chromatography (SEC). At each stage, eluted fractions and flow-through from the chromatographic columns were assayed for complement inhibitory activity, and the active fractions were further fractionated. First, protein extract was fractionated by anion exchange chromatography using a MonoQ 5/50 GL column (GE), washed with 10 column volumes (CV) 25 mM Na2HPO4/NaH2PO4, pH 7.0, and eluted by a 0–0.5 M NaCl gradient over 30 CV in 500 μL fractions. The flow-through was then acidified by addition of 1 μL 10 M HCl and injected onto a Dynamax 300-A□ C8 column (Rainin). The sample was eluted with a 0–80% ACN gradient in 0.1% TFA over 40 min. Aliquots were lyophilised and resuspended in 500 μL PBS. The active fraction was incubated at 21°C for 1 h with an equal volume of 3.4 M (NH4)2SO4, pH 7.0, centrifuged (22,000 x g, 10 min) and topped up to 0.95 mL with 1.7 M (NH4)2SO4, 100 mM Na2HPO4/NaH2PO4, pH 7.0. The sample was loaded onto a 1 mL HiTrap Butyl HP column (GE), and washed with 5 CV of 1.7 M (NH4)2SO4, 100 mM Na2HPO4/NaH2PO4, pH 7.0. Elution was carried out by a 1.7-0.0 M (NH4)2SO4 gradient over 15 CV in 1 mL fractions. All fractions were buffer exchanged to PBS and concentrated.

### Identification of novel tick inhibitors

Identified protein fractions with complement-inhibitory abilities were digested by Trypsin and analysed by LC-MS/MS. Samples were topped up to 50 μL with 50 mM TEAB, pH 8.5, reduced with 20 mM TCEP (21 °C, 30 min), alkylated with 50 mM chloroacetamide in the dark (21 °C, 30 min), digested with 0.5 μg of trypsin (37 °C, 16 h), then quenched with 1 μL formic acid. Digested peptides were analysed by LC-MS/MS over a 30 min gradient using LTQ XL-Orbitrap (Thermo Scientific) at the Central Proteomics Facility (http://www.proteomics.ox.ac.uk, Sir William Dunn School of Pathology, Oxford). Data were analysed using the central proteomics facilities pipeline (CPFP) (26) and peptides were identified by searching against the *R, pulchellus* sialome cDNA database (25) and an updated *R. pulchellus* sialome cDNA database from raw sequence data (Sequence Read Archive, NCBI, Accession no.: PRJNA170743) with Mascot (Matrix Science). Hits were assessed for the presence of a signal peptide with the SignalP 4.1 Server (27) (CBS, DTU), sequence homology to known protein sequences by blastp (NCBI), and structural homology to known protein structures by Fold and Function Assignment Service (FFAS) (28).

### *R. pulchellus* sialome cDNA database assembly

We downloaded female and male R. pulchellus sequencing data (100bp paired-end Illumina HiSeq 2000 reads) as published in (13). from the Sequence Read Archive (Accession ids SRX160117 and SRX160070, respectively). We de novo assembled female (51Mio read pairs) and male (70Mio read pairs) data independently using Bridger version (29) with default parameters. Raw reads were then mapped to the assembled female/male CDNA using NextGenMap 0.4.12 [(30), enforcing a minimum 95% sequence identify (-i parameter) and sorted read alignments were inspected in the IGV genome browser (31) for QC purposes.

### Expression and purification of recombinant proteins

#### Insect cell expression

Codon-optimized GeneArt strings were cloned into a modified pExpreS2-2 vector (ExpreS2ion Biotechnologies, Denmark) with an N-terminal His6 tag (pMJ41). The purified plasmid was transformed into S2 cells grown in EX-CELL 420 (Sigma) with 25 μL ExpreS2 Insect-TR 5X (ExpreS2ion Biotechnologies). Selection for stable cell lines (4 mg/mL geneticin (ThermoFisher)) and expansion were carried out according to the manufacturer’s instructions. **E. coli expression.** GeneArt strings were cloned into pETM-14 and transformed into T7 SHuffle cells (CirpT1-4) or BL21 (DE3) cells (C5_MG4), both cell types NEB. Protein expression was carried out in 2x YT broth (with 50 μg/mL kanamycin). Cells were induced with 1 mM IPTG. The cultures were centrifuged (3,220 x g, 10 min) and the cell pellets resuspended and lysed in PBS containing 1 mg/mL DNase and 1 mg/mL lysozyme by homogenization. Expressed proteins were subsequently purified by Ni-chelate chromatography (Qiagen, 5 ml column) and SEC (S75, 16 60, GE) in PBS.

#### Complement inhibition assays

Red blood cell hemolysis assays and complement ELISAs were carried out as described previously (9). In brief, **haemolysis assay** was performed with sheep red blood cells (TCS Biosciences) sensitized with Anti-Sheep Red Blood Cell Stroma antibody (cat. no. S1389, Sigma-Aldrich). 50 μl cells (5 x 108 cells/ml) were incubated in an equal volume of diluted serum (1 hour, 37 °C, shaking). Cells were pelleted and haemolysis was quantified at A405 nm of supernatant. Cells with serum only used for normalization (100% activity). Final serum dilutions used: 1/80 (human), 1/40 (rabbit), 1/160 (rat), 1/40 (pig) and 1/640 (guinea pig). Human serum was from healthy volunteers (prepared as described (9)), pig serum was a kind gift from Tom E. Mollnes (Oslo University Hospital, Norway), rat and guinea pig serum were from Complement Technology (USA) and rabbit serum was from Pal Freeze (USA). **Complement inhibition ELISAs** were performed using a Wieslab complement system screen (Euro Diagnostica, Sweden) following the manufacturer’s instructions, with sample added prior to serum. Contents were mixed by shaking at 600 rpm for 30 s.

#### Pull-down Assay

0.1 mg/mL of purified protein was immobilised on Pierce NHS-activated magnetic beads (ThermoFisher) following the manufacturers’ instructions. The beads were incubated with 10 mM EDTA and 50 μL serum (21°C, 30 min). The beads were washed thrice with 1 mL PBS + 0.05% Tween20, once with 100 μL PBS, and boiled in 50 μL SDS-PAGE loading buffer. The eluted proteins were separated on an SDS polyacrylamide gel and observed by Coomassie staining, silver staining, or Western blotting. For blotting the SDS-PAGE separated proteins were transferred to a PVDF membrane (Amersham Hybond P0.2 PVDF, 55 GE) by semi-wet transfer (BioRad) and blocked for 1 h with 2% milk. Primary antibody (α-C5: 1:80,000, Complement Technology, USA). Secondary antibody (α-Goat HRP, Promega, 1:10,000). The blot was developed using ECL Western Blotting Substrate (Promega) and imaged using Amersham Hyperfilm ECL (GE).

#### Purification of serum C5 and C5-inhibitor complexes

C5 and C5-inhibitor complexes were purified essentially as described (9). In brief, pre-cleared serum was incubated with His-tagged OmCI, and the C5-OmCI complex was purified by Ni-chelate and anion chromatography (Mono Q 10/100 GL column, GE). A two-fold molar excess was added of CirpT or RaCI inhibitors, or EcuFab (a custom-made Fab fragment prepared following the manufacturer’s framework (Ab00296-10.6, Absolute Antibody, UK) which includes the VL and VH sequences of Eculizumab (European Patent Office: EP0758904 A1). Following this SEC (S20010/30 HR column, GE) was used to remove excess inhibitors purify the final complexes (in PBS). SEC-MALS was performed as described (9).

### Surface Plasmon Resonance (SPR)

SPR experiments were performed using a Biacore T200 (GE). C5 was coupled to CM5 chips by standard amine coupling. CIRpT1-4 was flown over the surface in 0.01 M HEPES pH 7.4, 0.15 M NaCl, 3 mM EDTA, 0.005% v/v Surfactant P20, at a flow rate of 30 μL/min. The strong interaction was not sufficiently disrupted by either high/low salt (0-3 M NaCl) or extreme pH (range 2-8 tried) and extended dissociation time (1 h) was therefore used between successive injections. Fits were performed to control (blank channel)-subtracted traces. Data was fitted using a 1:1 Langmuir with mass transfer model. To calculate the affinity of CIRpT1-4 for C5, a series of injections at concentrations spanning ~3 nM to 2 μM were fit using the BiaEvaluation software.

### Cryo-electron microscopy, image processing, model building and refinement

4 μL of C5-OmCI-RaCI-CirpT1 in PBS (0.3 mg/ml) was applied to freshly glow-discharged (20 s, 15 mA) carbon-coated 200-mesh Cu grids (Quantifoil, R1.2/1.3). Following incubation for 10 s, excess solution was removed by blotting with a filter paper for 3 s at 100% humidity at 4□°C and plunge frozen in liquid ethane using a Vitrobot Mark IV (FEI). Data were collected on a Titan Krios G3 (FEI) operating in counting mode at 300 kV with a GIF energy filter (Gatan) and K2 Summit detector (Gatan). 4440 movies were collected at a sampling of 0.822 Å/pixel, dose rate of 6 e^−^/Å/s over an 8 s exposure for a total dose of 48 e^−^/Å^2^ over 20 fractions. Initial motion correction and dose-weighting were performed with SIMPLE-unblur (32) and contrast transfer functions (CTFs) of the summed micrographs were calculated using CTFFIND4 (33). Dose-weighted micrographs were subjected to picking using SIMPLE (32) fed with the known crystal structure of C5-OmCI-RaCI-complex (pdbid: 5hce). All subsequent processing was carried out using Relion 3.0-beta-2 (34). Movies were reprocessed using built-in MOTIONCOR2, with 5×5 patches and dose-weighting. Picked particles were extracted in a 288 x 288 Å box, totalling 502,640 particles. Reference-free 2D classification was performed and the highest resolution classes selected, leaving 35, 707 particles. 3D classification was then carried out using a low resolution *ab initio* model as a reference and the highest resolution class (118, 634 particles) was subjected to masked auto-refinement. Following Bayesian polishing and CTF refinement, gold standard Fourier shell correlations using the 0.143 criterion led to global resolution estimates of 3.5 Å. Post-processing was carried out using a soft mask and a B-factor of −106 Å^2^ was applied. Local resolution estimations were calculated within Relion 3.0. A model of C5-OmCI-RaCI1-complex (pdbid: 5hce) was fit into the map using the program COOT (33) and refined using the Real-Space Refinement module of Phenix (35). Volumes and coordinates have been deposited in the PDB with the ID 6rqj and the EMDB with ID: EMD-4983. See Data Table 1 for cryo-EM data collection, refinement and validation statistics.

### Crystallization, X-ray data collection, and structure determination

CirpT1 was co-purified with the C5_MG4 domain by SEC (S75 10/30, GE) in PBS and concentrated to 21 mg/ml. The protein complex was with an equal volume of mother liquor containing in 0.02 M Na2PO4/K2PO4, 20 % w/v PEG3350, and crystallized in 200 nL drops by vapor diffusion method at 21 °C. Crystals were cryoprotected in mother liquor supplemented with 30% glycerol and flash frozen in liquid N2. Data were collected on beamline I03 at the Diamond Light Source (Harwell, UK), wavelength: 0.9762 Å, as specified in Table 2. The structure of CirpT1-C5_MG4 was solved by molecular replacement using MolRep within CCP4 (36) with the structures of C5-OmCI-RaCI (PDB ID: 5HCC (9)). The structure of CirpT1 was manually built into difference density and the model subjected to multiple rounds of manual rebuilding in Coot (37) and refinement in Phenix (35). The structure of the complex is characterized by the statistics shown in Table 2 with 0 % Ramachandran outliers and 96.05 % of residues lying in the favorable regions of the Ramachandran plot. Structure factors and coordinates have been deposited in the PDB with the ID 6rpt. Interactions between CirpT1 and C5_MG4 have been predicted by PISA (38). Protein structure figures for both EM and X-ray structures were prepared using Pymol Version 2.0 (Schrödinger, LLC) and ChimeraX (39).

## Acknowledgements

We acknowledge Diamond Light Source and the staff of beamline I03 for access under proposal MX18069. We thank M. Slovak (Institute of Zoology, Bratislava, Slovakia) for providing salivary glands. We thank E. Johnson & A. Costin of the Central Oxford Structural Microscopy and Imaging Centre for assistance with data collection. H. Elmlund (Monash) is thanked for assistance with access to SIMPLE code ahead of release. The Central Oxford Structural Microscopy and Imaging Centre is supported by the Wellcome Trust (201536). M.P.R was financially supported by grants from the Wihuri foundation and the Finnish Cultural foundation. Staff and experimental costs in S.M.L. lab were supported by a Wellcome Investigator Award (100298) and an MRC programme grant (M011984).

## Author contributions

M.P.R.: Designed and performed experiments. Protein purification, characterisation of protein complexes, CryoEM grid optimization, X-ray crystallography, structure determination and analysis. Wrote paper with S.J. and S.M.L.

S.J.: Designed, supervised and performed experiments. Characterisation of protein complexes. CryoEM data analysis, structure determination and analysis. Wrote manuscript with M.P.R. and S.M.L.

M.J.: Designed, supervised and performed experiments. Fractionation of salivary gland proteins, identification of tick inhibitor. Strain and plasmid construction, protein purification, complement activation assays.

T.T.: Performed experiments. Fractionation of salivary gland proteins, identified tick inhibitor. Strain and plasmid construction, protein purification, complement activation assays.

T.M.: Performed experiments. Strain and plasmid construction. Protein binding studies.

N.T.: Performed experiments. Strain and plasmid construction. Protein purification.

N.P.: Performed experiments. Produced *R. pulchellus* sialome cDNA database.

J.D.: Performed experiments. CryoEM grid optimisation and data collection.

S.M.L.: Designed, supervised and performed experiments. CryoEM data optimisation and collection, data and structure analysis. Wrote paper and prepared figures with M.P.R. and S.J.

## Data deposition

X-ray coordinates and data have been deposited into the PDB with ID: 6rpt. The EM coordinates and volumes have been deposited into the PDB with ID: 6rqj and the EMDB with ID: EMD-4983.

